# Two new “Incertae sedis” syllids (Annelida: Syllidae) from Brazilian oceanic islands

**DOI:** 10.1101/2022.09.30.510401

**Authors:** Rodolfo Leandro Nascimento, Marcelo Veronesi Fukuda, Paulo Cesar De Paiva

## Abstract

Oceanic islands present very interesting environments, known by possessing relatively distinct fauna and flora. However, taxonomic accounts from Brazilian oceanic islands focused on important groups, such as the family Syllidae, began to be published only in recent years. In this paper we provide descriptions and illustrations of two new species, *Brevicirrosyllis* sp. nov. from Trindade Islands and *Westheidesyllis* sp. nov. from Rocas Atoll, two incertae sedis genera previously included in the Eusyllinae subfamily. We also provide updated identification keys for both genera.

## Introduction

The family Syllidae Grube (1850) is one of the most complex and species-rich groups of annelids, with 79 genera, comprising about 1100 valid species (San Martin & Aguado 2014; Pamungkas *et al.* 2019; Martin *et al.* 2021). Currently, the family is divided into five subfamilies: Anoplosyllinae Aguado & San Martín, 2009, Autolytinae Langerhans 1879, Eusyllinae Malaquin, 1893, Exogoninae Langerhans, 1879, and Syllinae Grube, 1850, in addition to some *“Incertae sedis*” genera (Aguado *et al.* 2012; San Martin & Aguado, 2014).

Aguado *et al.* (2012) combined morphological and molecular information to elucidate the relationships within Syllidae, and found “Eusyllinae” as paraphyletic, while also finding a monophyletic group within “Eusyllinae”; thus, the authors proposed a reorganization, conserving the name and the rank of subfamily. In this process, some genera were considered as ‘*Incertae sedis*’ – excluded from the new configuration of the Eusyllinae –, many of them for which no molecular information was available at the time; this is the case, among others, of *Brevicirrosyllis* San Martin, López & Aguado, 2009 and *Westheidesyllis* San Martin, López & Aguado, 2009 (Aguado *et al.* 2012; 2015).

Both *Brevicirrosyllis* and *Westheidesyllis* count with only one official record each in Brazilian waters, as *Brevicirrosyllis* cf. *mariae* (San Martin & Hutchings, 2006), recorded from Southeastern Brazil (Fukuda *et al.* 2015), and *Westheidesyllis gesae* (Perkins, 1981) (as *Pionosyllis gesae*), from the Rocas Atoll (Paiva *et al.* 2007) – the former, however, lacking descriptions and information on deposited material. Here we describe two new species, *Brevicirrosyllis* **sp. nov.**, from Trindade island, the sixth known species of the genus, and *Westheidesyllis* **sp. nov.**, the first species of the genus reported with glands, described from the Rocas Atoll.

## Material and Methods

Specimens were collected in two oceanic islands from Northeastern Brazil. *Westheidesyllis* **sp. nov.** was found in the Rocas Atoll (3°51’S 33°40’W), the only atoll in the South Atlantic Ocean, at 260 km off the coast of Natal, Rio Grande do Norte, Northeastern Brazil; specimens were fixed in formalin 10% and, later, preserved in ethanol 70%. The specimens of *Brevicirrosyllis* **sp. nov.** were found in the Trindade Island (20°30’S 29°20’W), collected through the ‘ProTrindade Marine Invertebrate Project’ (‘*Pro Trindade*’), focused on the fauna of the Trindade and Martin Vaz Archipelago, at 1140 km off the coast of Vitória, Espírito Santo, Southeastern Brazil; specimens from this project were both fixed and preserved in ethanol 70%.

Morphological traits were analysed and measured under a Zeiss Stemi SV11 stereomicroscope and Zeiss Axio Lab A1 microscope. In addition, some specimens were examined using scanning electron microscopy (SEM). For SEM, specimens were first dehydrated in a graded series of increasing concentrations of ethanol (92–100%), critical point-dried, coated with ~35 nm of gold, and examined and photographed at the Laboratório de Imagem e Microscopia Óptica e Eletrônica (LABIM–UFRJ). Line drawings were done from slide-mounted specimens with the aid of a drawing tube. The length of specimens was measured from the tip of palps to the tip of pygidium, excluding anal cirri; width was measured at proventricular level, excluding parapodia. Type material and other examined specimens are deposited at the Museu Nacional, Universidade Federal do Rio de Janeiro (MNRJP), Brazil, and at the Museu de Zoologia, Universidade de São Paulo (MZUSP), Brazil.

## Results

> Class Polychaeta Grube, 1850
>
> Subclass Errantia Audouin & Milne Edwards, 1832 Order Phyllodocida Dales, 1962
>
> Family Syllidae Grube, 1850
>
> *Incertae sedis*
>
> Genus ***Brevicirrosyllis*** San Martín, López & Aguado, 2009
>
> Type species: *Pionosyllis weismanni* Langerhans, 1879, designated by San Martín *et al.* (2009).

### Diagnose

Small to medium sized syllids, usually slender, without ciliary bands. Palps triangular, distally rounded, fused at bases and diverging towards tips. Prostomium frequently with two pairs of eyes and two anterior eyespots, some species without eyes, and three antennae, median antenna inserted posteriorly on prostomium. Peristomium usually distinct, with 2 pairs of peristomial cirri. Antennae, peristomial and dorsal cirri of chaetiger 1 long and slender; from chaetiger 2 onwards, dorsal cirri short, usually ovate to exogonid-like. Ventral cirri digitiform. Compound chaetae as hemigomph or heterogomph falcigers, with blades bidentate to subbidentate and spinulated, with short spines on margin; dorsoventral gradation in length present. Dorsal simple chaetae present from at least chaetiger 2; ventral simple chaetae present on posterior body, bidentate, sometimes with hood covering subdistal tooth. Proventricle and pharynx about same size; pharyngeal tooth located anteriorly, near anterior margin. Reproduction by epigamy (cf. San Martin *et al.* 2009).

### Remarks

*Pionosyllis* Malmgren, 1867 was redefined by San Martín *et al.* (2009), with the proposition of some new genera – *Brevicirrosyllis* among them – based on coherent groups previously encompassed within that genus. In that work, the authors provided an identification key for the species of *Brevicirrosyllis*, but the information about the morphology of the dorsal simple chaetae of *B. mayteae* (San Martin & Hutchings, 2006) and *B. ancori* (San Martin & Hutchings, 2006) were exchanged from one to the other: in the original description and illustrations the former species has the dorsal simple chaetae pin-shaped, while the latter has the dorsal simple chaetae truncate (San Martin & Hutchings 2006). We update this identification key and insert the new species described herein.

> ***Brevicirrosyllis* sp. nov.**

### Diagnosis

*Brevicirrosyllis* without dorsal cirri on second parapodium, parapodial glands absent, palps similar in length to prostomium, median antenna more than four times longer than palps, dorsal peristomial cirri longer than body width.

### Type material

**Holotype. Brazil**, Trindade Island, Enseada da cachoeira (−20.2357136, −28.6968285), 18 m depth (MZUSP 2027), coll.04. July. 2012. **Paratype**. Trindade Island, Ilha da Racha (−20.5083781, −29.4587883), 21 m depth: 1 specimen (MZUSP 2267), coll.16. July. 2013.

### Additional material examined

*Brevicirrosyllis ancori* (San Martín & Hutchings, 2006). Australia, Queensland, Great Barrier Reef, Outer Younge Reef (−14.6149208, 145.5871437), rock covered with coralline algae and encrusting sponges, 9 m: 1 spec. (holotype, AM W29244), coll. P. Hutchings, 21 Jan 1977, det. G. San Martín, 15 Nov 2004; same locality, rock with *Lithothamnion* and *Halimeda*, 30 m: 4 specs (AM W28962), coll. P. Hutchings, 24 Jan 1977, det. G. San Martín, 2003.

### Description

Medium to long-sized body, slender, longest specimen examined 7 mm long, 0.17 mm wide, with 47 chaetigers; body without pigmentation in specimen preserved in ethanol (Fig. 1A). Palps triangular, distally tapering, fused only at bases, about same length of prostomium; prostomium subpentagonal, with a pair of eyes at anterior 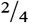 of its length, and a pair of eyespots on anterior margin; lateral antennae inserted slightly anteriorly to pair of eyes, about same length of palps; median antenna inserted posteriorly to eyes, on middle of prostomium or slightly posteriorly, almost four times longer than lateral ones (Fig. 1A). Peristomium distinct, shorter than subsequent segments; dorsal peristomial cirri about same length of palps and prostomium together, longer than body width and the lateral antennae, but shorter than dorsal cirri from chaetiger 1; ventral peristomial cirri about ⅓ length of dorsal ones (Fig. 1A). Dorsal cirri from chaetiger 1 longer than remaining ones, with almost half length of median antenna; dorsal cirri absent from chaetiger 2; remaining dorsal cirri digitiform to distally slightly tapered, without internal glands, longer than parapodial lobes but shorter than width of respective chaetiger (Fig. 1A). Ventral cirri digitiform, shorter than parapodial lobes. Parapodial lobes conical. Anterior body parapodia with 5–4 falcigers each, 4–3 falcigers on midbody and 3 falcigers on each posterior body parapodium; shafts of falcigers smooth, thicker ventralwards; blades bidentate, distal tooth larger than subdistal one throughout; on each parapodium, dorsalmost blade elongate, subdistally faintly sinuous (Fig. 1B); blades with short and thin spines on margin, with smooth connective joining blade and shafts on anterior and midbody parapodia; blades with dorsoventral gradation in length, about 27–10 μm on anterior body (Fig. 1B), 29–8 μm on midbody (Fig. 1C) and 15–7 μm on posterior body parapodia (Fig. 1E). Dorsal simple chaetae present from chaetiger 10–11, truncated, with few spines laterally, becoming slightly thicker towards posterior body (Fig. 1F, G, H). Ventral simple chaetae not observed. One acicula per parapodium throughout, almost bent at right angle, with irregular, tapering tip (Fig. 1D). Pharynx through 2.5–3 segments; with a conical to rhomboidal pharyngeal tooth located on anterior rim (Fig. 1A). Proventricle through 2.5 segments, with 32–30 muscle cell rows.

**Fig. 1.**
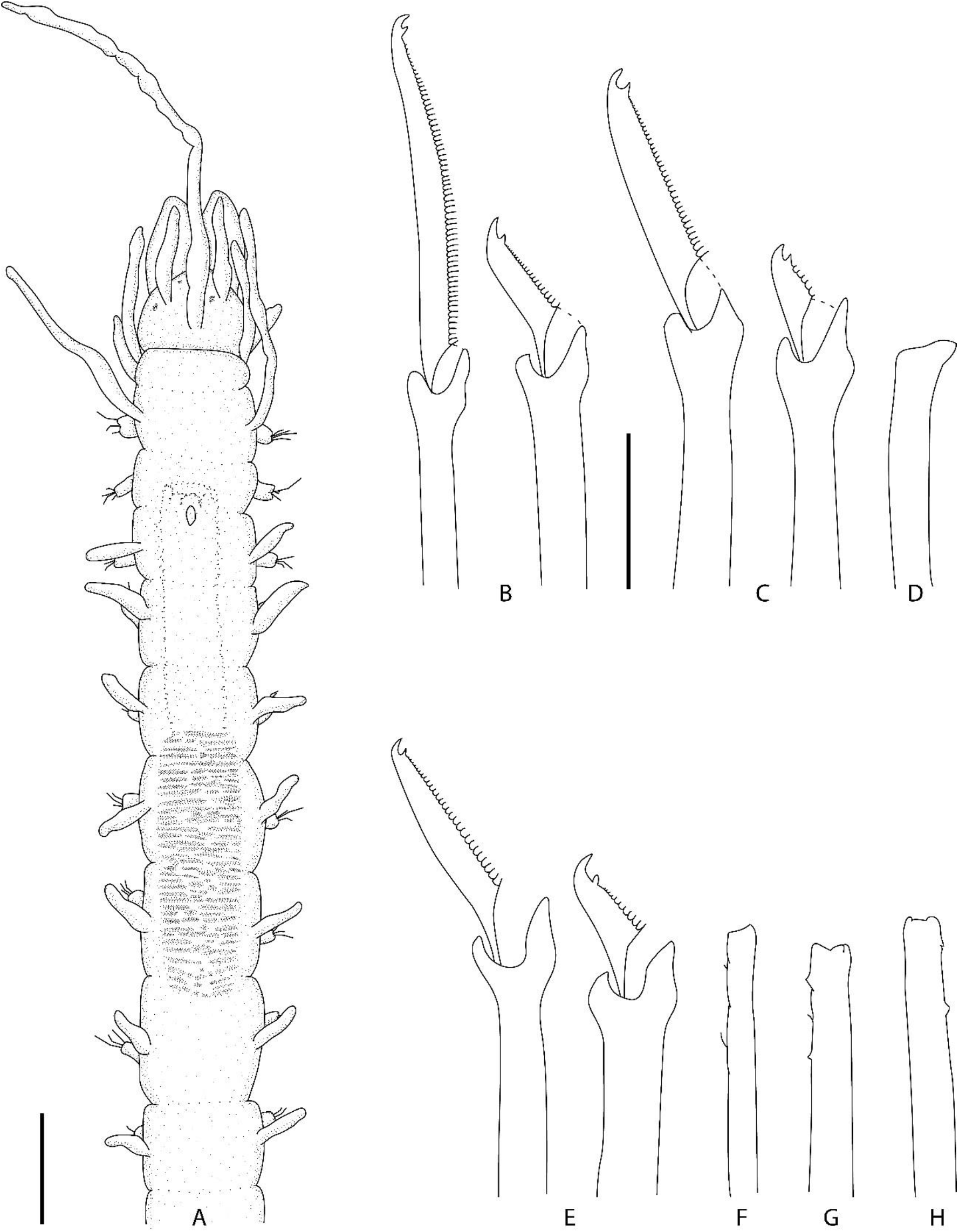
*Brevicirrosyllis* sp. nov. **A**. Anterior body. **B, C, E**. Falciger chaetae of anterior, mid- and posterior body, respectively. **D**. Aciculae. **F–H**. Dorsal simple chaeta of anterior, mid- and posterior body, respectively. Scale bar: **A**, 0. 17 mm; **B–H**, 10 μm.

### Remarks

*Brevicirrosyllis* **sp. nov.** is characterized by having palps with about same length of prostomium; median antenna more than four times longer than palps; dorsal peristomial cirri with about same length of palps and prostomium together, longer than body width; dorsal cirri of chaetiger 1 longer than remaining, about half length of median antenna or twice longer than width of corresponding segment; and dorsal cirri of remaining chaetigers shorter, digitiform to distally slightly tapered, lacking internal glands.

*Brevicirrosyllis ancori* described from Queensland, Australia, in the Pacific Ocean, is the most similar species to *B.***sp. nov.**, sharing the overall body morphology and shape of compound chaetae. Conversely, *B. ancori* differs by having palps about 1½ longer than prostomium, median antenna shorter, with only about twice the length of palps, dorsal peristomial cirri about same size of palps, about same length of body width, and by having parapodial glands. Moreover, San Martin & Hutchings (2006) described some variation in *B. ancori:* on some specimens, two pairs of eyes may be present; dorsal cirri on chaetiger 1 may be longer (cf. San Martín & Hutchings, 2006, Fig. 56A–B) and parapodial glands larger than in the holotype. Even compared with these specimens that varied from the type-series of *B. ancori, B.***sp. nov.** can be easily differentiated.

### Type locality

Trindade island, Espírito Santo, Brazil.

### Distribution

South Atlantic Ocean, Trindade island.

Identification key to the currently known species of ***Brevicirrosyllis*** (adapted from San Martin *et al.* 2009)

1. Ventral simple chaetae without hood, about same width of falciger shafts, with both teeth similar in size. Compound chaetae heterogomph ................ ***Brevicirrosyllis gorringensis*** Hartmann-Schröder, 1977.

– Ventral simple chaetae with or without hood, wider than falciger shafts, with subdistal tooth longer than distal one. Compound chaetae hemigomph ................. 2

2. Dorsal cirri with fibrillar inclusions ................. 3

– Dorsal cirri without fibrillar inclusions ............... 4

3. Dorsal cirri on chaetiger 2 present ... ***Brevicirrosyllis weismanni*** (Langerhans, 1879)

– Dorsal cirri on chaetiger 2 absent ***Brevicirrosyllis mariae*** (San Martin & Hutchings, 2006)

4. Dorsal simple chaeta pin-shaped ***Brevicirrosyllis mayteae*** (San Martin & Hutchings, 2006)

– Dorsal simple chaeta truncated ................5

5. Palps longer than prostomium; median antenna twice length of palps; dorsal peristomial cirri about same length of body width; parapodial glands present .................. ***Brevicirrosyllis ancori*** (San Martin & Hutchings, 2006)

– Palps with same length of prostomium; median antenna longer than above, more than four times length of palps; dorsal peristomial cirri longer than body width; parapodial glands absent ............ ***Brevicirrosyllis* sp. nov.**

> **Genus *Westheidesyllis* San Martin, López & Aguado, 2009**

### Type species

*Eusyllis heterocirrata* Hartmann-Schröder, 1959, designated by San Martín *et al.* (2009). **Diagnosis (Emended)**. Small-sized, fragile bodies, easily loosing antennae and cirri. A transversal band of cilia may be present on prostomium, peristomium and segments. Palps subtriangular, free from each other for most of their length, fused only at bases; prostomium oval to subpentagonal, with lateral antennae inserted near anterior rim, median antenna inserted posteriorly to lateral ones; eyes present or absent, sometimes only a pair; some species with pair of anterior eyespots. Nuchal organs as transversal ciliated grooves between prostomium and peristomium. Peristomium distinct, with two pairs of peristomial cirri. Dorsal cirri alternating long cirri, more than twice longer than body width at corresponding segment, and short cirri, with length up to half width of corresponding segment. Ventral cirri digitiform, inserted distally on parapodial lobes. Parapodial glands occasionally present at the bases of parapodial lobes. Falcigers with homogomph articulation; blades short, bidentate, spinulated, with short spines. Dorsal simple chaetae from anterior to midbody posteriorwards. Ventral simple chaetae not known. Aciculae distally inflated, laterally expanded or knobbed. Pharynx longer or about same size as proventricle, with anterior tooth (cf. San Martin *et al.* 2009).

### Remarks

Since its proposal, the genus *Westheidesyllis* counted with only three species: *W. corallicola* (Ding & Westheide, 1997), *W. gesae* (Perkins, 1981) and *W. heterocirrata* (Hartmann-Schröder, 1959). *Westheidesyllis gesae* was recorded for Brazilian waters (as *Pionosyllis gesae)*, specifically for the Rocas Atoll (Paiva *et al.* 2007), however, this record lacks a description and details on deposited material. Here we describe *Westheidesyllis* **sp. nov.** also from the Rocas Atoll, the first species of the genus reported as presenting glands, which lead us to amend the genus to conform this character.

> ***Westheidesyllis* sp. nov.**

### Material examined

#### Holotype

Rocas Atoll (−3.8805091, −33.8780718), 1 m depth, on coralline sand (MNRJP XXXX), coll. 16 Oct 2000. **Paratypes.** Rocas Atoll (−3.8805091, −33.8780718), 1 m depth, on coralline sand 4 specimens (MNRJP XXXX), coll. 16 Oct 2000.

Rocas Atoll (−3.8805091,-33.8780718), 1 m depth, on coralline sand: 135 specimens, coll. 16 Oct 2000; Piscina das Âncoras (−3.8638266, −33.8263897), 1 m depth, on coralline sand: 57 specimens (four mounted for SEM), coll. 16 Oct 2000; “along of the Rais”, 1m depth, on coralline sand: 6 specimens, coll. 23 Oct 2000.

### Additional material examined

*Westheidesyllis gesae* (Perkins, 1981). United States, Florida, St. Lucie County, Hutchinson Island (27.3567, −80.2217), 10.9 m: 1 spec. (holotype, USNM 60456), coll.

Gallagher, Boyle & Whiting, 12 Mar 1976, det. T.H. Perkins; same locality (27.3689, 80.2294), 9.7 m: 1 spec. (paratype, USNM 60458), coll. Gallagher, Futch & Jaap, 29 Jul 1973, det. T.H. Perkins (1 spec.); same locality (27.3564, −80.2233), 11.5 m: 2 specs (paratypes, USNM 60459), coll. Gallagher & Hollinger, 14 Mar 1972, det. T.H. Perkins. *Westheidesyllis heterocirrata* (Hartmann-Schröder, 1959). El Salvador, Estero Jaltepeque, La Herradura, sand, infralittoral: 1 spec. (holotype, HMZ P-14579), 1955.

### Description

Small-sized, slender bodies, longest specimen 2.6 mm long, 0.25 mm wide, with 32 chaetigers; specimens preserved in ethanol without pigmentation. Palps subtriangular, basally juxtaposed for ~1/4 their length, distally rounded, slightly shorter than prostomium (Figs. 2A; 3A; 4A; 5A, C, D). Prostomium ovate to subpentagonal; eyes absent; lateral antennae inserted close to anterior margin of prostomium about half length of median one; median antenna inserted on midline of prostomium, almost four times longer than palps and prostomium (Figs. 3; 5A–D). Two large ciliated nuchal organs between prostomium and peristomium (Fig. 5A, B). Peristomium distinct, shorter than subsequent segments; dorsal peristomial cirri about same length or slightly shorter than median antenna (Fig. 3); ventral peristomial cirri almost half length of dorsal ones.

**Fig. 2.**
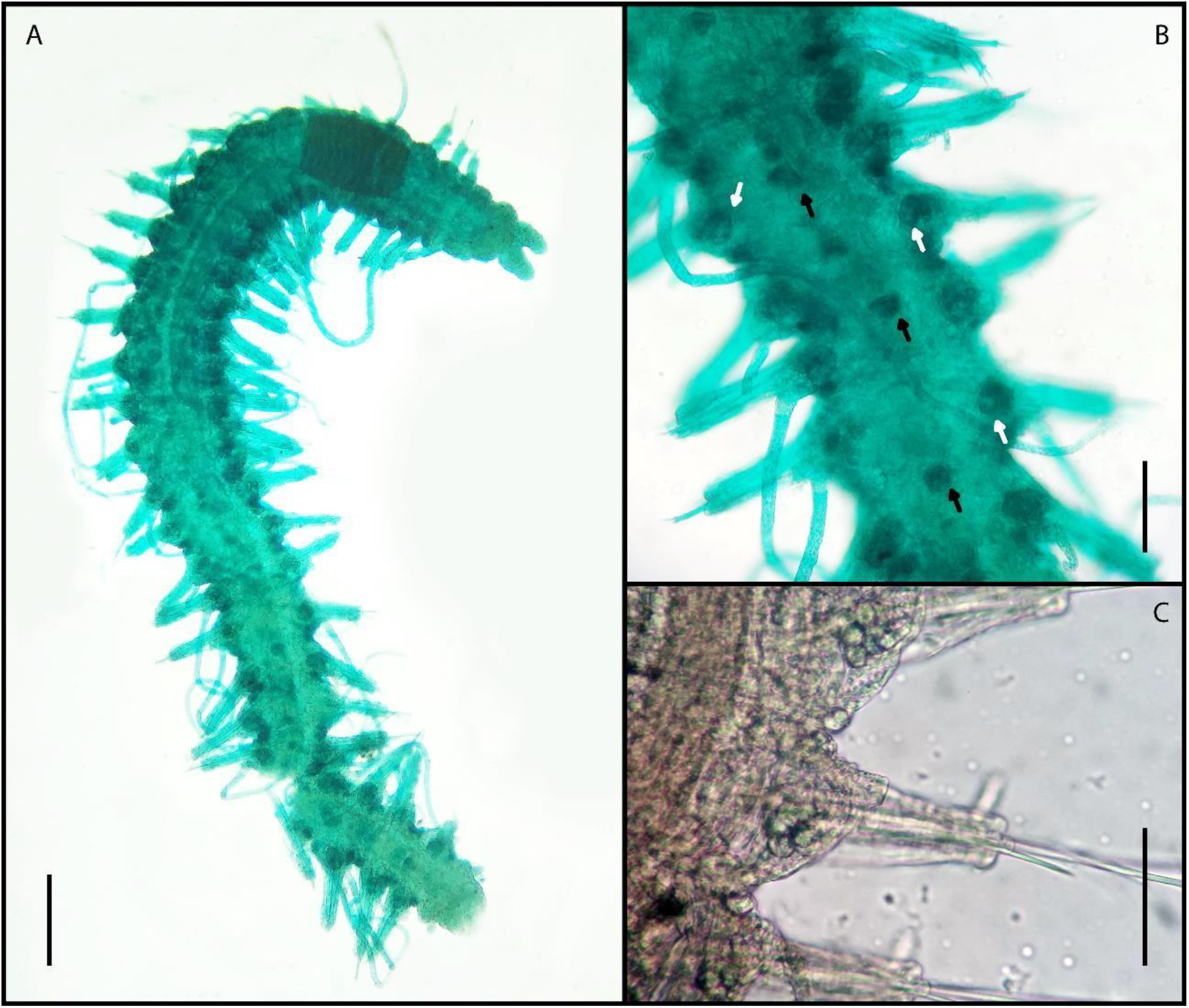
*Westheidesyllis* sp. nov. **A**. Whole body, dorsal view. **B**. Midbody of a methyl green stained specimen, white arrows showing parapodial glands, black arrows showing glands on the digestive tube. **C**. Midbody parapodial glands on a regular (not stained) specimen. Scale bar: **A**, 0. 22 mm; **B**, 0.15 mm; **C**, 0.1 μm.

**Fig. 3.**
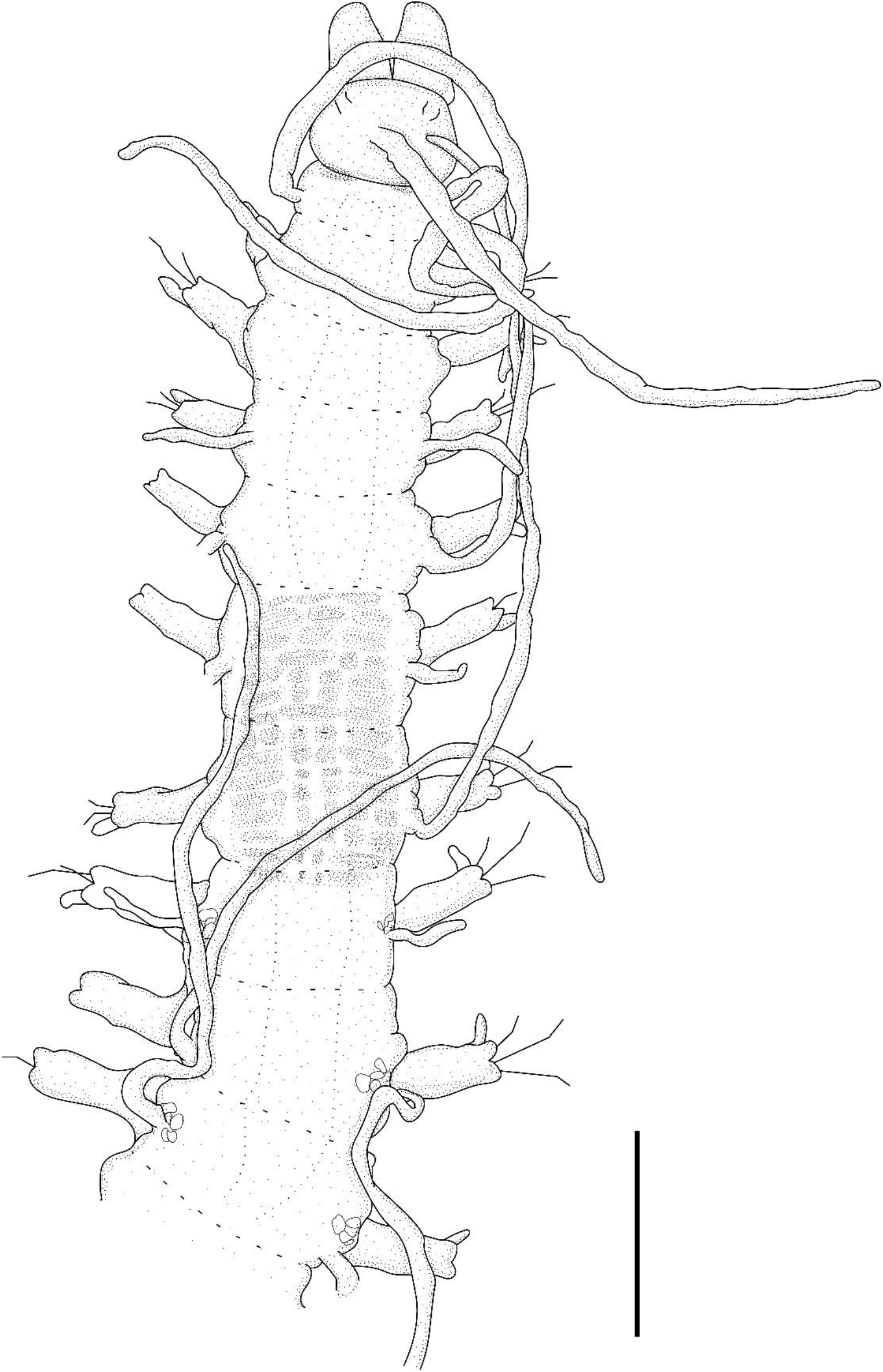
*Westheidesyllis* sp. nov. **A**. Anterior body, dorsal view. Scale bar: 0.22 mm.

**Fig. 4.**
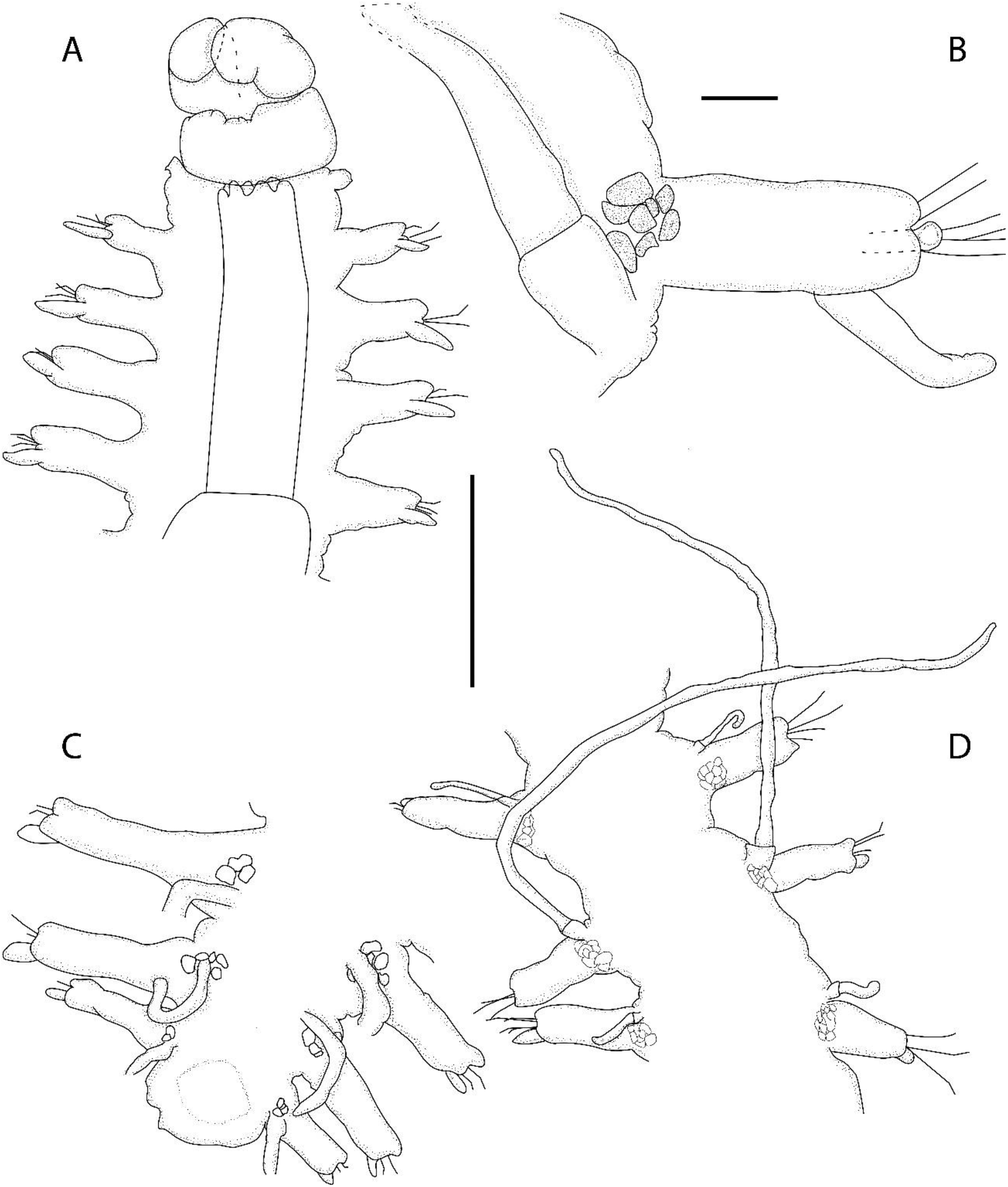
*Westheidesyllis* sp. nov**. A**. Anterior body, ventral view. **B**. Midbody parapodia, with dorsal cirrus and parapodial glands, dorso-lateral view. **C–D**. Midbody and posterior end, showing segments, parapodial lobes and glands, dorsal cirri, dorsal view. Scale bars: **A**, **C**, **D**, 0.2 mm; **B**, 15 μm.

**Fig. 5.**
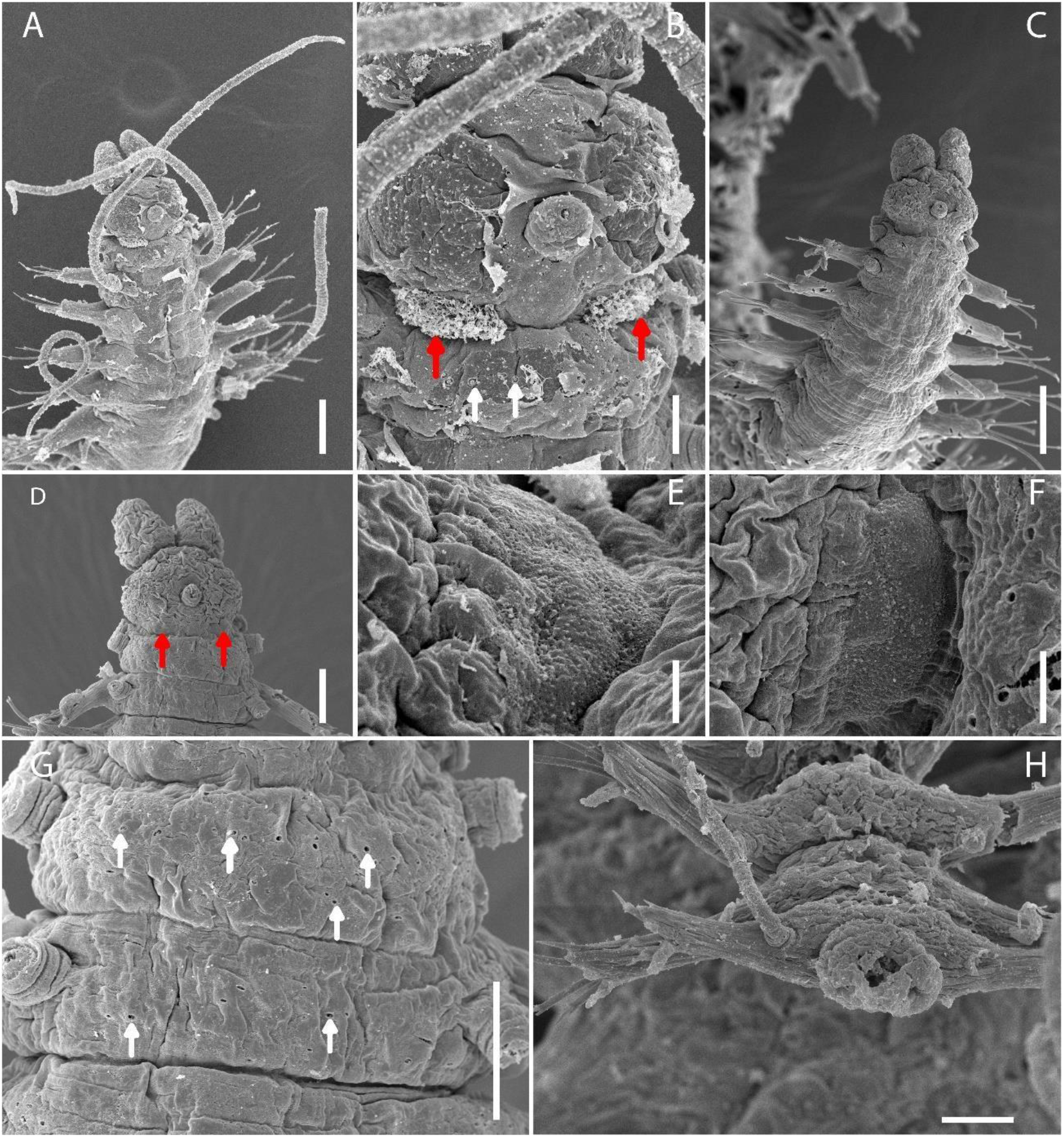
*Westheidesyllis* sp. nov. SEM. **A**. Anterior body, dorsal view. **B**. Details of prostomium and peristomium, dorsal view. **C**. Anterior body of specimen with retracted nuchal organs, dorsal view. **D.** Anterior end, dorsal view. **E–F**. Details of retracted ciliated nuchal organs; **G.** Anterior segments showing details of ciliary pits, dorsal view. **H**. Posterior end, dorsal view. Red arrows pointing ciliated nuchal organs, white arrows pointing ciliary pits. Scale bars: **A**, **C**, 50 μm; **B**, **G**, 10 μm; **D**, **H**, 20 μm; **E**, 2 μm; **F**, 5 μm.

Ciliated pits transversally arranged on midline of peristomium and segments, to at least chaetiger 15 (Fig. 5B, G). Dorsal cirri alternating in length, on chaetiger 1 about four times longer than width of segment (Fig. 3); on chaetiger 2 absent; on chaetigers 3, 5 and 7 shorter than width of corresponding segment; on chaetigers 4, 6, 8 and 9 three to four times longer than width of corresponding segment (Fig. 3); from chaetiger 10 onwards, dorsal cirri with regular alternation in length, short cirri shorter than corresponding segment, long cirri three to five times longer than corresponding segment, (Fig. 4D).

Antennae, peristomial and dorsal cirri with cirrophores (Figs. 3A; 4B, D). Ventral cirri digitiform, shorter than parapodial lobes, inserted distally, extending beyond parapodial lobes, shorter towards posterior body (Figs. 4A, B; 5H). Parapodial lobes elongated, rectangular, slightly bilobed (Fig. 4B); parapodial glands presents after proventricle, on the bases of parapodial lobes, with rounded to subpentagonal granules (Figs. 2A–C; 4B– D). Parapodia with three falcigers throughout; shafts of falcigers smooth, homogomph, with irregular, usually quadrilobate acute tips (Fig. 7F); blades bidentate, with teeth about same size or distal tooth slightly larger throughout; blades spinulated, with short and thin spines (Figs. 6A–C; 7A, B, F, J); blades varying in length on dorsalmost, intermediate and ventralmost chaetae, with 6 μm, 12 μm and 8 μm on anterior parapodia (Figs. 6A; 7A,B); 7 μm, 13 μm and 10 μm long on midbody (Figs. 6B; 7E, F); and 5 μm, 12 μm and 9 μm on posterior body (Figs. 6C; 7H–J). Dorsal simple chaetae present from chaetiger 3–4, tapering distally, with rounded tip, subdistally spinulated on anterior body (Figs. 6D; 7C, D), becoming slightly sigmoid towards posterior body (Figs. 6E, F; 7G, K). One acicula per parapodium throughout, distally inflated, hollow (Fig. 6G), with tip protruding from parapodial lobe (Fig. 4B). Pharynx through about 4 segments (Figs. 2A; 3), with conical to rhomboidal pharyngeal tooth located on anterior rim, surrounded by 10 soft papillae; proventricle through 2.5 segments, with 14–15 muscle cell rows (Fig. 3).

**Fig. 6.**
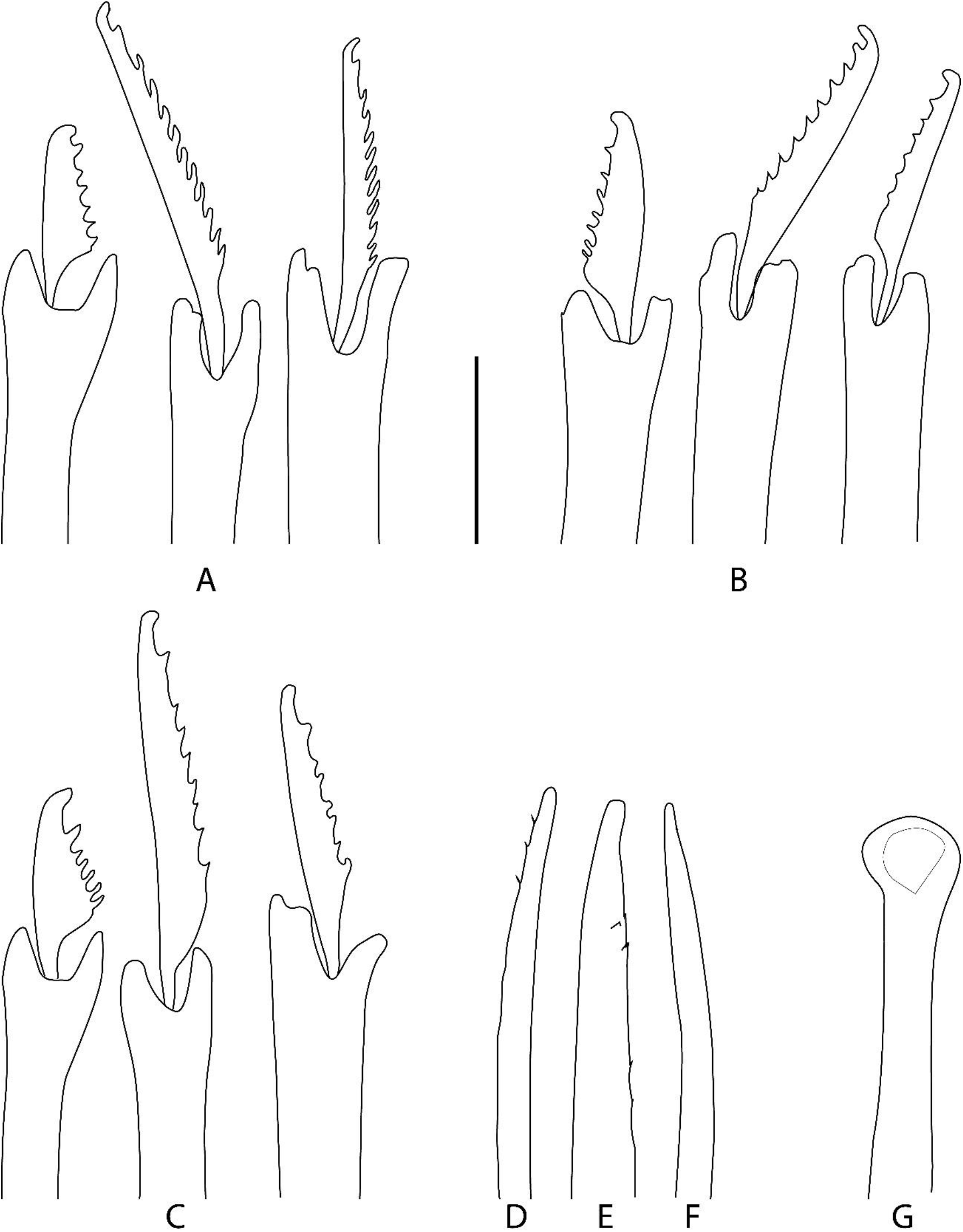
*Westheidesyllis* sp. nov. **A–C**. Falcigers chaetae of anterior, mid- and posterior body; **D–F**. Dorsal simple chaetae of anterior, mid- and posterior body. **G**. Acicula. Scalebar: 6 μm.

**Fig. 7.**
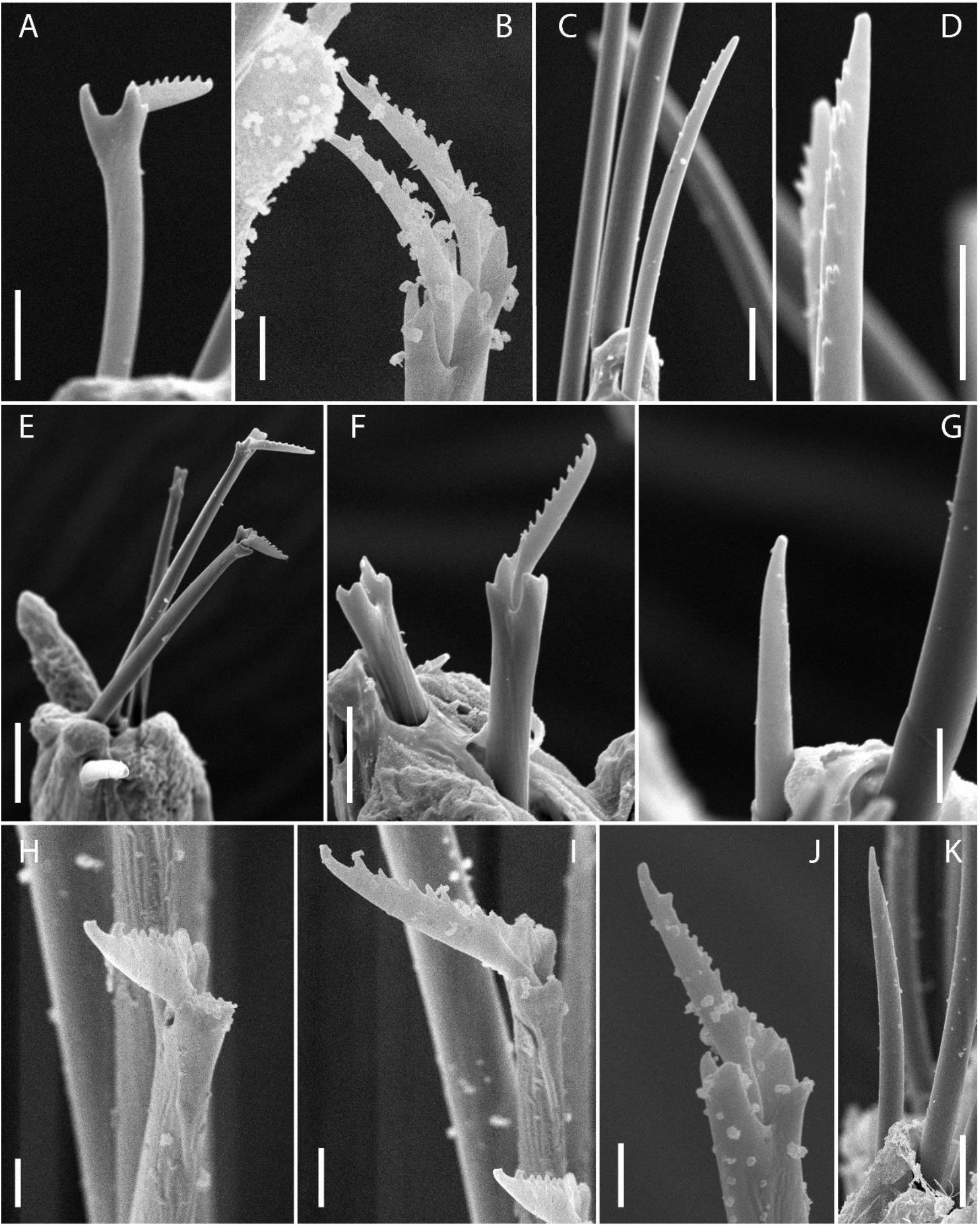
*Westheidesyllis* sp. nov. SEM. **A**. Falcigers chaetae of anterior body. **C–D**. Dorsal simple chaeta of anterior body and details of its tip, respectively. **E–F**. Falcigers of midbody. **G**. Dorsal simple chaeta of midbody. **H–J**. Falcigers chaetae of posterior body. **K.** Dorsal simple chaeta of posterior body. Scale bars: **A**, **C**, **F**, 5 μm; **B**, **D**, **G**, **I**, **J**, **K**, 2 μm; **E**, **H**, 10 μm.

Pygidium rounded (Figs 4C; 5H), with pair of cirri about same length of long posterior body dorsal cirri.

### Remarks

None of the specimens of *Westheidesyllis* **sp. nov.** examined herein showed cilia at the bases of the dorsal cirri or the transversal ciliary bands on the segments throughout, as mentioned in other species of the genus. Nonetheless, under SEM, it was possible to observe a set of pits, of which, generally, these cilia are projected: at the bases of the dorsal cirri, almost above the parapodial glands and arranged transversely, more or less in line, on each anterior segment and peristomium.

*Westheidesyllis* **sp. nov.** resembles *W. corallicola* (Ding & Westheide, 1997), described from Hainan Island, South China, and later found in Australia (New South Wales and Lizard Island), all records in the Pacific Ocean. Members of both species lack eyes, also sharing the overall body morphology and similar compound chaeta. *Westheidesyllis* **sp. nov.** lacks eyespots, have median antenna inserted medially on prostomium, aciculae distally hollow, with tips protruding from parapodial lobes, and proventricle extending for 2.5 segments, besides the internal glands on the bases of parapodia. Conversely *W. corallicola* has eyespots, median antenna inserted posteriorly on prostomium, aciculae distally knobbed but not hollow nor protruding from parapodial lobes (Ding & Westheide, 1997, Fig. 6D, E, I), and proventricle extending for about 1.5 segment (Ding & Westheide, 1997, Fig. 6A), and lacking internal glands (Ding & Westheide 1997; San Martin & Hutchings 2006). Furthermore, specimens of *Westheidesyllis* **sp. nov.** showed no signs of cilia nor the ciliary pits indicating a similar ciliation pattern to that found in *W. corallicola*, regarding the tufts dorsally and ventrally located close to the bases of parapodia and on the pygidium (Ding & Westheide, 1997).

As mentioned above, *Westheidesyllis* **sp. nov.** is the only known species of the genus where glands have been observed. The presence of glands, specially associated to the parapodia, on interstitial species in soft-bottom substrates is commonly reported (Worsaae *et al.*, 2021). The parapodial glands in *Westheidesyllis* **sp. nov.** are best observed after Methyl green staining (Fig. 2A, B), but they can be relatively easily visualized without the aid of this technique (Fig. 2C).

#### Type locality

Rocas Atoll.

#### Distribution

Atlantic Ocean: Rocas Atoll, Brazil.

**Key to the current known species of *Westheidesyllis* (adapted from San Martin *et al.* 2009)**

1. Eyes absent, but anterior eyespots may be present ................2

– Eyes and eyespots present ..................................3

2. Without eyespots; parapodial glands present; aciculae distally hollow, with tips protruding from parapodial lobes ................. *Westheidesyllis* **sp. nov.**

– With eyespots; parapodial glands absent; aciculae distally knobbed, not protruding from parapodial lobes ...... *W. coralicolla*

3. Transversal ciliated bands on prostomium, peristomium and segments; blades of falcigers with long and thin spines ............... *W. gesae*

– Transversal ciliated bands absent, or not as above; blades of falcigers with spines coarser than above ........................ *W. heteroccirata*

## Discussion

Two species of the genus *Westheidesyllis*, *W. gesae*, described from Florida and with reports from the Atlantic coast of the United States, Gulf of Mexico and the Caribbean, (Read & Fauchald, 2021), and *W. heterocirrata*, described from and only known to occur in El Salvador, in the Pacific Ocean (Read & Fauchald, 2021), are morphologically very similar to each other. *Westheidesyllis gease* has anterior and midbody falciger blades with long and thin spines, ciliation on the prostomium and as transversal ciliary bands in each segment, and proventricle extending for about three segments, with ca. 23 muscle-cell rows. On the other hand, *W. heterocirrata* presents falciger blades with spines relatively thicker, proventricle extending for about two segments, with 14 muscle-cell rows, and does not have transversal ciliary bands in the segments.

The clear identification of ciliation patterns can be very tricky without proper fixation methods and examination under SEM (San Martín & Aguado, 2012), which difficult identifications in genera for which this character is important, as is the case of *Westheidesyllis.* Illustrating the issue, ciliation in some paratypes of *W. gease* could not be visualized under optical microscopy (MVF, pers. obs.); accordingly, Salcedo *et al.* (2016) found that the transverse ciliary bands might not be present in some specimens of *W. gesae* from the Mexican Pacific. On the other hand, clear tufts of cilia could be observed on the base of cirrophores in the holotype of *W. heterocirrata* (MVF, pers. obs.), although this character was not mentioned in the original description (Hartmann-Schröder 1959). Therefore, we recommend that revisions of the species within this genus, ideally with SEM, should be performed, in order to clarify the status of these taxa.

